# Predicting off-target effects for end-to-end CRISPR guide design

**DOI:** 10.1101/078253

**Authors:** Jennifer Listgarten, Michael Weinstein, Melih Elibol, Luong Hoang, John Doench, Nicolo Fusi

## Abstract

The CRISPR-Cas9 system provides unprecedented genome editing capabilities. However, off-target effects lead to sub-optimal usage and additionally are a bottleneck in development of therapeutic uses. Herein, we introduce the first machine learning-based approach to this problem, yielding a state-of-the-art predictive model for CRISPR-Cas9 off-target effects which outperforms all other guide design services. Our approach, Elevation, consists of two inter-related machine learning models—one for scoring individual guide-target pairs and another which aggregates guide-target scores into a single, overall guide summary score. Through systematic investigation, we demonstrate that Elevation performs substantially better than competing approaches on both of these tasks. Additionally, we are the first to systematically evaluate approaches on the guide summary score problem; we show that the most widely-used method (and one re-implemented by several other servers) performs no better than random at times, whereas Elevation consistently outperformed it, sometimes by an order of magnitude. In our analyses, we also introduce a method to balance errors on truly active guides with those which are truly inactive, encapsulating a range of practical use cases, thereby showing that Elevation is consistently superior across the entire range. We thus contribute a new evaluation metric for benchmarking off-target modeling. Finally, because of the large computational demands of our tasks, we have developed a cloud-based service for end-to-end guide design which incorporates our previously reported on-target model, Azimuth, as well as our new off-target model, Elevation.

## Introduction

Although the CRISPR-Cas9 system is routinely used, potentially avoidable off-target effects are likely reducing its effectiveness. The best way to mitigate off-target effects is to know when and where they occur and then design guides to avoid them while balancing for on-target efficiency.^1,2^ Such a balance may differ for different tasks. For example, the generation of cellular and animal models, or therapeutic uses of CRISPR-Cas9, will in general be far less tolerant of off-target effects than genome-wide screens wherein redundancy of targeting can be used to average out off-target effects. Nevertheless, reduction of off-target effects is desirable in all applications.

While GUIDE-seq^3^, IDLV^4^, Digenome-seq^5,6^ and other laboratory-based assays^1^ can be used to quantify off-target effects, scaling these assays to all guides genome-wide, against all genome-wide potential off-targets, is not practically feasible owing to cost, labor and availably of general-purpose assays.^1^ In contrast, as we show herein, machine learning-based predictive modelling can leverage a small number of such data to learn statistical regularities of guide-target sequence pairs that cause off-target effects, as well as their aggregate effect on a cell. Such modelling thus enables cheap and rapid *in silico* screening of off-target effects at a genome-wide level for guides never before assayed.^1,2,7^

There are two main uses cases for off-target predictive-modelling. The first is to understand how active a given guide–off-target region is likely to be, which we refer to as *guide-target scoring*. This is useful if one is concerned about a particular region of the genome, such as accidentally knocking out a tumor-suppressor gene when trying to make an edit to disable an HIV entry receptor. The second use case is to obtain an overall *summary score* of all off-target region activity for a given guide, so as to for a rank-order of good potential guides. This is useful to ensure overall precise activity for both therapeutic and screening applications. These two tasks are interrelated in that the summarization task requires accurate individual guide-target pair scoring. Moreover, to perform the summarization task requires selecting which potential off-target regions to score for a given guide, which we refer to as *search and filter.* Therefore, one can break down the off-target predictive modelling problem into three main tasks: given a guide to evaluate for off-target activity one needs to (i) *search and filter* genome-wide for potential targets, for example, those regions of the genome matching the guide up to *n* number of mismatches; (ii) *score* each potential target for activity from step i; (iii) *aggregate* the scores from step ii into a single off-target potential with which to assess the guide.

A number of solutions have been presented for the first task of search and filter, including Cas-OFFinder^8^, CRISPOR^9^, CHOP-CHOP^10^, e-CRISPR^11^, CRISPR-DO^12^ and CROP-IT^13^, which differ in the algorithms used to search as well as the completeness of the search. Completeness is dictated by options such as maximum number of mismatches, allowed protospacer adjacent motifs (PAMs) and the search algorithm used. For infrastructural efficiencies and ease of integration with our cloud service, we created our own system to perform search and filtering, but using the same parameters as in ref.^1^ The second and third steps, of scoring and aggregation, have been explored considerably less than the search and filter step and are the focus of this work. A list of available guide design services that perform one or both of these steps is shown in Table 1. The only existing tools that return aggregation scores are the MIT web server^14^, CRISPOR^9^ and CRISPR-DO^12^, the latter two which re-implement the MIT web server rules. CHOP-CHOP^10^ counts the number of potential off-targets without scoring them; CROP-IT^13^ uses a hand-crafted series of rules and has been shown to be substantially outperformed by the MIT web server.^9^ Additionally, the CFD method^1^ has been shown to outperform the MIT web server on guide-target pair scoring^9^, but the CFD web server does not perform aggregation.

**Table 1.** Summary of CRISPR guide design services which include off-target scoring.

| Shorthand | On-target scoring | Off-target scoring | Off-target aggregator | On-target interface | Off-target interface |
| --- | --- | --- | --- | --- | --- |
| Elevation (this paper) & Azimuth<sup>1</sup> | new machine-learning based models | new machine-learning based models | Yes, machine-learning based | human genic targets pre-computed; cloud API for re-use in code and Excel; source code | human genic targets pre-computed web site; source code |
| MIT server<sup>14</sup> | new hand-crafted rules | new hand-crafted rules | Yes, hand-crafted rules | web site | web site |
| CRISPR-DO<sup>12</sup> | re-uses rules from Xu *et al.*<sup>16</sup> | re-uses rules from MIT server<sup>14</sup> | Yes, as in MIT server | web site; source code | web site; source code |
| CRISPOR<sup>9</sup> | re-uses rules from multiple papers<sup>1,16–21</sup> | re-uses rules from MIT server<sup>14</sup> | Yes, as in MIT server | web site; source code | web site; source code |
| Broad GPP<sup>1</sup> | new machine-learning based models | newly developed rules based on data | No | web site; source code | web site; source code |
| E-CRISPR<sup>11</sup> | new hand-crafted rules, and rules from ref. <sup>12,16,18</sup> | new hand-crafted rules | Yes, hand-crafted rules | web site | web site |
| CHOP-CHOP<sup>10</sup> | re-uses rules from Xu *et al.*<sup>16</sup> by default, and ref. <sup>1,17,18</sup> | counts # of off-targets but does not score them | No | web site | web site |

Although new CRISPR systems are being developed which may increase specificity (e.g., CPF115), these systems are still in their infancy, and Cas9 remains the workhorse endonuclease of choice. Moreover, only Cas9 has enough data to perform modelling at this point in time; hence our focus on Cas9 herein.

For each of guide-target scoring and guide summary scoring we developed machine learning approaches which substantially improve upon the state-of-the-art for the respective task, as demonstrated through our experiments. Together, we call our end-to-end modelling of off-targets, Elevation, which complements our on-target model, Azimuth. A schematic of our approach is shown in Figure 1.

**Figure 1.**
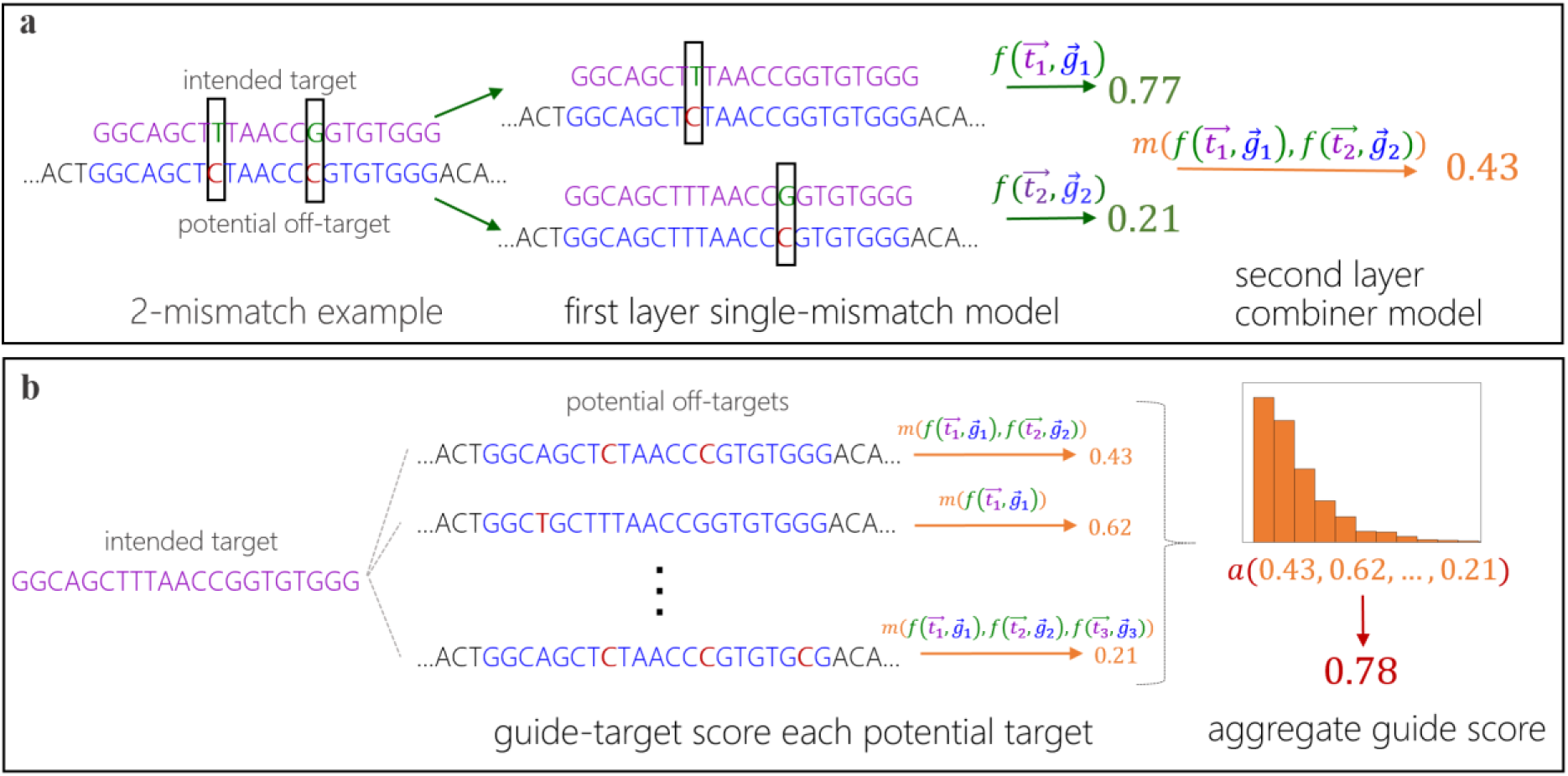
Schematic of Elevation off-target predictive modelling. **(a)** An example of how to score a guide-target pair with two mismatches. First the guide-target pair is broken down into two single-mismatch pseudo-pairs, each of which is scored with the first layer model. Then these scores are combined with the second-layer model, yielding a single guide-target score. **(b)** An example how to aggregate the set of guide-target scores for a single guide into one summary off-target score for a guide. The aggregator model computes statistics of the input distribution of guide-target scores as features for the model.

For the guide-target scoring task we developed a two-layer stacked regression model wherein the first layer (based on gradient-boosted regression trees) learns to predict the off-target activity of a *single-mismatch* (including both RNA:DNA mismatches as well as non-NGG PAM) guide-target pair while the second-layer model (based on an L1-penalized regression) learns how to combine predictions from the first-layer single-mismatch model for guide-target pairs with multiple mismatches into a single guide-target score. To apply the two-layer model to a multiple mismatch guide-target pair, one first decomposes each mismatch into a set of pseudo single-mismatch guide-target pairs, applies the first-layer model to each of these, then combines their outputs into a single score using the second-layer model. This procedure can be applied to a guide-target pair for any number of mismatches, although we have trained and evaluated on only up to six mismatches. Because of the combinatorial explosion of possible mismatch combinations, the relative amount of training data for the combiner task is extremely small. Consequently, we used a parsimonious model here—L1-penalized linear regression. Details and intuitions for development of the two-layer Elevation-score model are given in the Online Methods. Note that indels contribute to the off-target problem to a much lesser extent^3^, hence we have focused our modeling efforts on mismatches.

To build an aggregation predictive model which summarizes the individual guide-target scores in to a single number, we first compute a set of features for each guide based on the distribution of off-target scores for that guide (*e.g.*, the mean, median, percentiles, variance, sum of genic scores, fraction of genic scores over all potential target site scores)—see Methods for full description. We then use these features with gradient-boosted regression trees, which allows them to interact in the predictive model.

## Results

In this section, we first we evaluate guide-target pair prediction approaches, including our newly developed, Elevation-score, demonstrating that it yields state-of-the-art performance. Then we make use of Elevation-score to evaluate Elevation-aggregate with the two competing summarization approaches—the MIT web server, and CFD, where only the former has an accompanying web service which provides summary scores (we use scores computed from ref.^1^ to perform the CFD comparison).

### Individual guide-target pair off-target predictive modelling

We evaluated Elevation-score using two independent data sets generated from genome-wide unbiased assays—one based on GUIDE-Seq^3^, and the other an aggregated data set from Haeussler *et al.*^9^ Elevation-score outperformed all other models (CFD^1^, the current state-of-the-art, Hsu-Zhang^2^, and CCTop^7^) in predicting off-target activity (Figure 2). Note that for off-target prediction it is generally more damaging to mistake a truly active off-target site for an inactive one, rather than the other way around, because only the first type of error can disrupt the cell or confound experimental interpretation, while the second may only require designing another guide. Consequently, we chose an evaluation measure which accounts for this asymmetry—the weighted Spearman correlation, where each guide-target pair is weighted by an amount which is a (monotonic) function of its measured activity. Because the precise asymmetry is not *a priori* known and may vary for different applications, we varied the weight continuously between two extremes: from being directly proportional to the measured activity (such that false negatives effectively do not count relative to false positives), to a uniform weighting (*i.e*., yielding standard Spearman correlation). Note that we do not break down performance by number of mismatches because when using a Spearman rank correlation, one could do very well within each mismatch category, but terribly on the whole, and it is the whole that is what is of interest. Additionally, we do not break down analysis by mismatches as compared to non-NGG PAMs because the GUIDE-Seq and Haeussler data did not have examples of only non-NGG PAMs (rather they always also have a mismatch), making such an analysis impossible.

**Figure 2.**
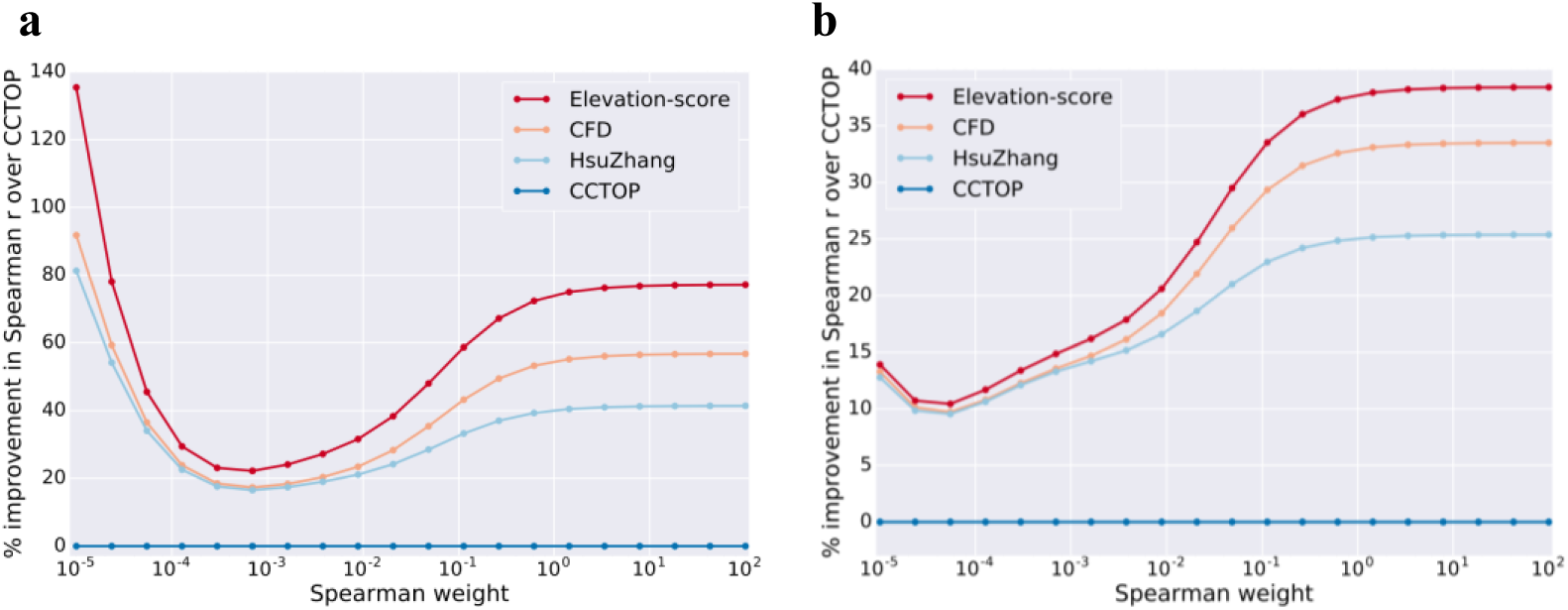
Guide-target pair scoring. Comparison of Elevation-score to other methods, evaluated using a weighted Spearman correlation between predictions and assay measurements. The horizontal axis shows different weights in the weighted Spearman—at the far left the weight is effectively proportional to the rank-normalized GUIDE-Seq counts/cutting frequency, while at the far right the weight is effectively uniform, yielding a traditional Spearman correlation. For ease of visualization, the vertical axis denotes the percent improvement of each model over CCTOP, which by design thus lies constant at zero. **(a)** CD33 and GUIDE-Seq data were used to train, while Hauessler *et al* data (after removing the GUIDE-Seq) were used to test. **(b)** the roles of the GUIDE-Seq and Haeussler data are reversed from **a**. The final Elevation-score model deployed in our cloud service uses the model trained on GUIDE-Seq data. Note that respectively only 0.12% and 0.51% of count values in GUIDE-Seq and Hauessler are non-zero, making the actual Spearman correlation difficult to interpret. For completeness, however, the right-most points correspond to a correlation of respectively 0.125, 0.111, 0.010 and 0.070 for Elevation, CFD, Hsu-Zhang and CCTOP (left plot) and 0.059, 0.057, 0.053 and 0.043 (right plot). The p-values for each correlation surpassed the floating point error and were reported as p=0.0 in all cases, demonstrating that despite the somewhat low correlations, a tremendous amount of signal is present.

For first-layer (single-mismatch) model features we used the position of the mismatch, the nucleotide identities of the mismatch, the joint position and identities of the mismatch in a single feature, and whether the mutation was a transition or transversion. The importance of these features is shown in Figure 3. It is interesting to note that using both the joint “position and mismatch nucleotide identity” features—those effectively used by CFD—are aided by additionally de-coupling into additional features of position and nucleotide identity, even though regression trees can in principle (with enough data) recover the joint features from the decoupled ones. Using only the CFD features in our model, or using classification instead of regression, or omitting the second-layer machine learning each caused the model to perform worse (Supplementary Figure 1). Feature importances for the second-layer (multiple-mismatch combiner) model show that the product and sum of the first layer single-mismatch predictions are driving the model, but that individual components of these need to be modulated by machine-learning for optimal performance.

**Figure 3.**
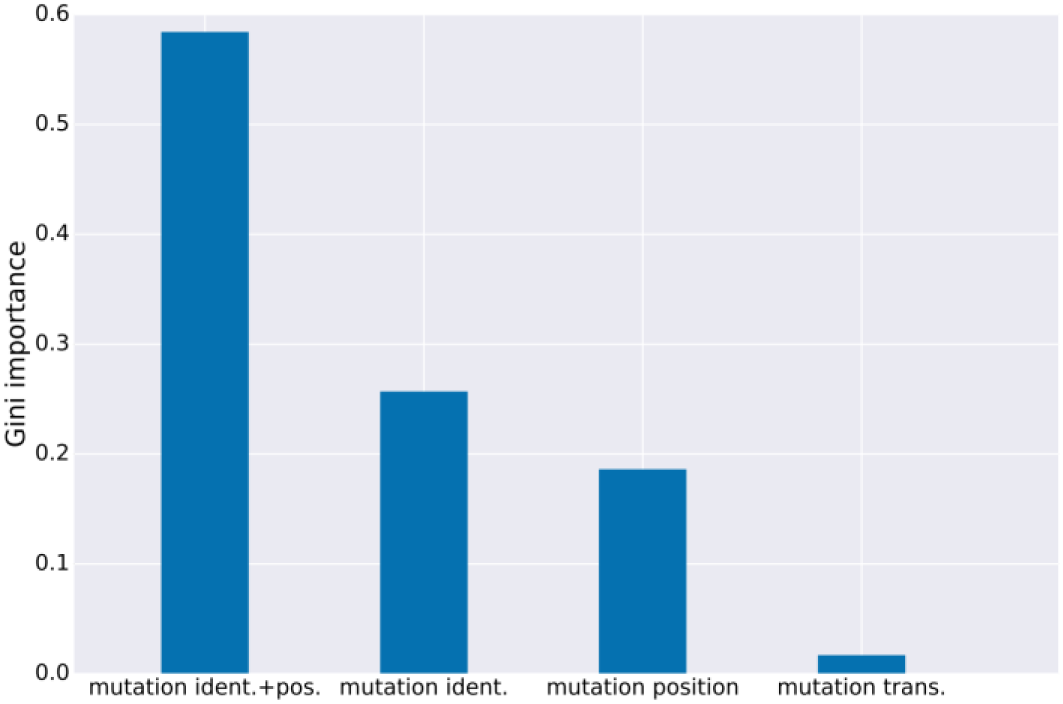
First-layer guide-target scoring feature importances. Average Gini importances for type of features in the first-layer single-mismatch boosted regression trees model (mutation nucleotide identities and position jointly; mutation identity; mutation position; mutation transversion *vs.* transition). The Gini importance refers to the decrease in mean-squared error (the criterion used to train each regression tree) when that feature is introduced as a node in the tree. This measure has a close, empirical correspondence with the importance of the feature that would be obtained with a permutation test, and can also be viewed as a relative decrease in entropy provided by splitting on that feature. This measure of importance does not convey whether having that feature makes a guide better or worse in the model because such a notion is impossible for regression trees in which the effect of one feature is dependent on the presence/absence of other features (*i.e.*, there are non-linear interactions between the features). This model was trained with CD33 single-mismatch and alternative PAM data.

### Aggregating individual off-target scores into a single guide summary score

The end task, of aggregation, requires obtaining a single off-target *summary score* for a guide given all its individual guide-target scores. This task is likely of most interest for guide design. To evaluate our approach on this task we made use of two data sets with guides targeting non-essential genes in viability screens, the Avana^1^ and Gecko^22^ libraries. Because each guide is designed to target one non-essential gene in these screens, the cell should be viable if no off-targets effects are present. However, to the extent that off-target effects are present, the cell is more likely to die, which is captured by the assay. Hence this kind of experiment serves as bronze standard for the combined task of scoring and aggregation, and also enables us to develop a machine learning approach to aggregation.

**Figure 4.**
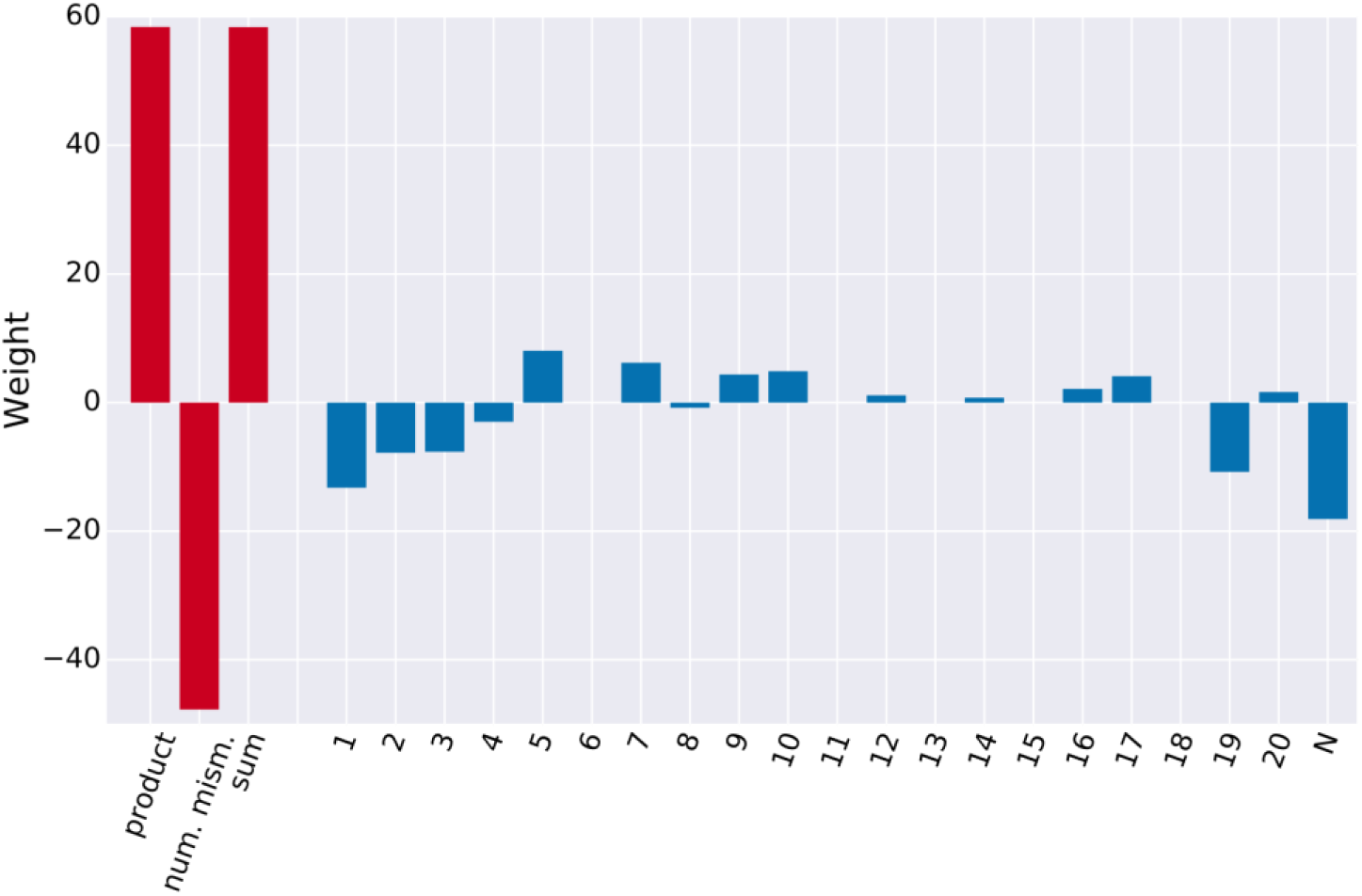
Second-layer guide-target scoring feature importances. Weights from second-layer predictive model in Elevation—an L1-regularized regression. This model uses (the log of) the single-mismatch predictions from the first layer as inputs (one for each position in the 20-mer guide and the N position of the PAM, using a 1.0 if no mismatch occurred at that position), shown in blue. The remaining inputs are the product of these non-logged values of these features, as well as their sum, and number of mismatches (and alternate PAM) in the guide, shown in red. The use of the product feature by itself, without the other 23 features would yield the model Elevation-naïve which performs worse. As a consequence, the other features in Elevation can be viewed as a correction of the assumptions in the development of Elevation-naïve, such as independence of mismatches and the regression yielding a true probability.

Aggregation of Elevation-score to obtain a summary score was the best-performing model; however, we found that adding CFD-based guide-target score features further boosted performance and so also included these in the final Elevation model. Elevation yielded up to a 6-fold improvement (and was never worse) in Spearman correlation over the best approach for this end-point task—the CFD website—and an even larger improvement over crispr.mit.edu^2^, the most widely-used guide design tool (Figure 5) which sometimes performed at random. The importance of each aggregation feature is shown in Figure 6. Note that in general, models with richer predictive modelling capacity are more difficult to interpret, so there is a tradeoff between model interpretability and predictive accuracy. We focused solely on accuracy as the intent was to provide the best tool possible for the guide design community.

**Figure 5.**
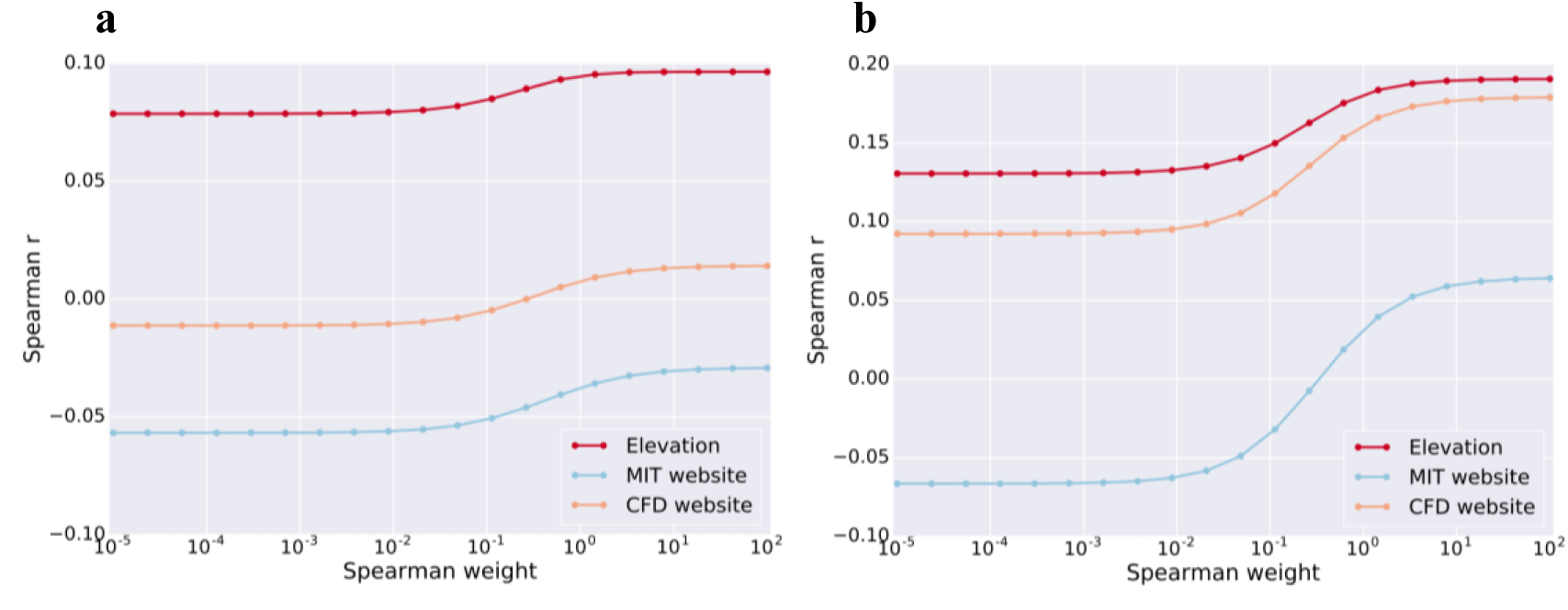
Joint scoring and aggregation on viability screens. Spearman correlation of Elevation to the crispr.mit.edu server. **(a)** Avana data was used to train and Gecko to test, **(b)** the reverse of **a**. Note that the MIT website yields correlation in the wrong direction for most settings in both data sets. The final Elevation model deployed in our cloud service uses the model trained on Avana.

**Figure 6.**
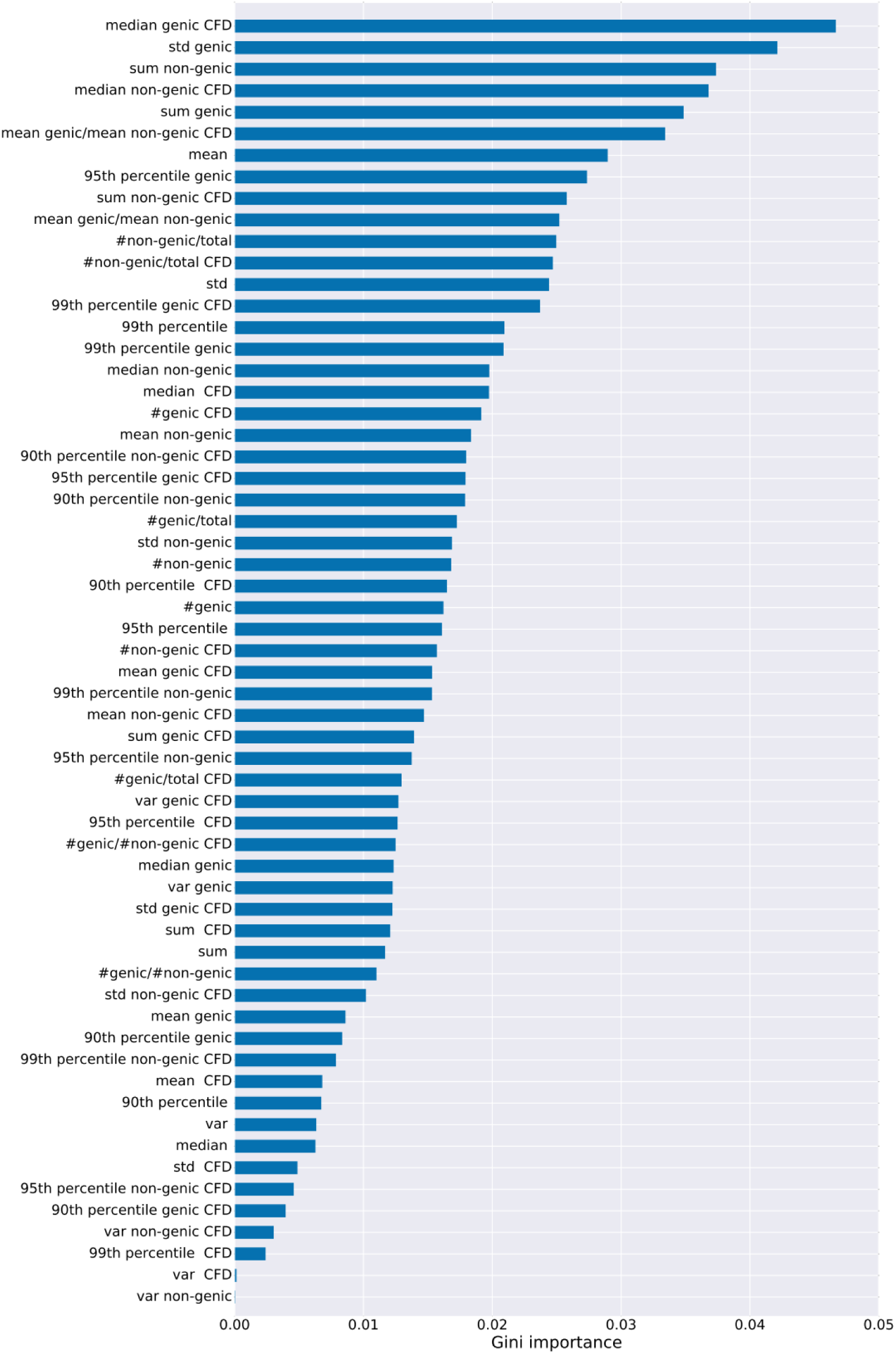
Aggregator feature importances. Weights from aggregator model in Elevation which uses Gradient Boosted regression trees. The features were: the mean; median, variance, standard deviation, 99^th^, 95^th^, 90^th^ percentiles, and sum of off-target scores. We compute these for each of: all off-targets, only genic off-targets, and only non-genic targets. Additionally, we compute these further features: sum of genic [non-genic] off-targets divided by total number of off-targets; fraction of targets that are genic; fraction that are non-genic; ratio of number of genic to non-genic targets; ratio of average genic to non-genic score. Those with a postfix of “CFD” are from that model, while all others are from Elevation-score. The Gini importance is described in Figure 3.

## Conclusion

We have introduced the first machine-learning based approach to predictive modelling of off-target effects for CRISPR-Cas9. Through systematic investigation, we demonstrated that our newly developed suite of models, Elevation, performs better for each of the two main off-target-related tasks in guide design. Additionally, we are the first to systematically evaluate available competing approaches on the task of summary scoring (aggregation), showing that the most widely-used method (and one re-implemented by several other tools) can perform no better than random at times, whereas Elevation consistently outperformed this approach by an order of magnitude. Additionally, we considered how to balance errors on truly active guides with those which are truly inactive, developing a new metric to do so, based on the weighted Spearman correlation. This type of evaluation encapsulates a range of practical use cases, and enabled us to show that Elevation is consistently superior across the entire range. We recommend that the community should use such metrics in future when comparing new and existing models for off-target modelling.

As data become available for a richer set of scenarios, including different endonucleases, different organisms and *in vitro* versus *in vivo*, we will update our models and tools accordingly. At present, there is not enough data to perform off-target modelling for more than CRISPR-Cas9.

Elevation-score and Elevation-aggregate, together referred to as Elevation, complement our on-target predictive model, Azimuth.^1^ Together, Azimuth and Elevation along with our cloud service and web front end, provide an integrated end-to-end guide design tool that enable users to more effectively deploy CRISPR-Cas9 for screening and therapeutics—one based on the state-of-the-art machine-learning based methods. In future work we will also more carefully investigate the issue of search and filter.

## Materials and Methods

### Data

To train our first-layer, single-mismatch model, we used CD33 data from Doench *et al.*^1^ where all single-mismatch mutations and alternate PAMs were introduced into the target DNA for 65 perfect-match guides that were effective at knockout. CD33-negative cells were isolated by flow cytometry so that their log-fold-change prior to CRISPR-Cas9 introduction could be measured by sequencing. After filtering as in Doench *et al.*, we retained 3,826 single-mismatch observations and 1,027 alternate PAM observations for a total of 4,853 guide-target training examples of which 2,273 were considered active by Doench *et al*. These data measure protein knockout efficiency rather than DNA cleavage. We refer to these data as the CD33 data.

To train our second-layer, multiple mismatch model, we used two unbiased/genome-wide multiple mismatch data sets. The first were GUIDE-Seq data^3^ comprising nine guides assessed for off-target cleavage activity. These guides yielded 354 active off-target sites (*i.e.*, non-zero counts) with up to six mismatches. Non-active sites were obtained from Doench *et al.* where Cas-OFFinder^8^ was used to identify all 294,310 sites with six or fewer mismatches. The second data comprised off-target data aggregated by Haeussler *et al.*^9^, after removing GUIDE-Seq data to make it independent from the previously mentioned data set. These data consisted of 122 active targets among 23,357 non-active potential targets. We applied a per-guide normalization (dividing by the total number of counts for a guide) to the GUIDE-Seq data to adjust for possible artifacts between guides, as done by Haeussler *et al.* We also set the minimum resulting value to 0.001, the estimated sensitivity of the assay^9^. Finally, we Box-Cox transformed^23^ both the GUIDE-Seq and Haeussler *et al.* activity layers.

Our goal was to build a tool for predicting knockout activity, rather than cleavage. However, given the paucity of either type of data and the fact that cleavage is required for knockdown, we chose to use both knockdown and cleavage data, as has been previously done^1^. Moreover, because our model generalizes well to independent off-target cleavage data, and does so better than other models, our model is likely to also prove useful for cleavage prediction.

To evaluate the aggregation of off-target effects we used to data sets arising from guides targeting non-essential genes in a viability screen. The first, from the Avana library^1^, used 4, 950 guides targeting 880 non-essential genes. The second, from the Gecko library^22^, used 4,697 guides targeting 837 non-essential genes).

### Predictive Modelling for Scoring Individual Guide-Target Pairs

Here we describe the *CFD* model^1^, what assumptions it makes, and then describe our model, Elevation-score, and how it relates conceptually to *CFD*. The predictive off-target model, *CFD*, first computes the observed frequency of guide-target pair activity for each single-mismatch type and alternative PAM in the CD33 data. *CFD* then combines these single-mismatch frequencies by multiplying them together for guide-target pairs with multiple mismatches. For example, if a guide-target pair had a A:G mismatch in position 3, a T:C mismatch in position 5 and a PAM of “CG” in the target region, then *CFD* would take the off-target score of this guide to be *CFD score* = *P*(*active* |*A*: *G*,3) × *P*(*active*|*T*:*C*,5) × *P* (*active*|*CG*), where each of the these terms is computed from observed frequencies in the CD33 training data (which contained only single-mismatch, or alternate PAMs, but never both).

#### CFD as Naïve Bayes

One can interpret the *CFD* algorithm in terms of a known classification model called Naïve Bayes^24^, as follows. First, denote *Y* = 1 to mean a guide-target pair was active, and *Y* = 0 to denote that the pair was not. Next, denote features such as T:C,5 as *X*_*i*_, where *i* simply indexes some enumeration of these features (*i.e*., a one-hot encoding). If that feature (mismatch) occurred, then *X*_*i*_ = 1, and if it didn’t occur then *X*_*i*_ = 0. Therefore, in the CD33 data (with only single mismatches/alternate PAM), a particular guide-target pair has only one *X*_*i*_ = 1 and all others have *X*_*i*_ = 0. In this notation one can re-write *CFD* as follows for one guide-target pair:

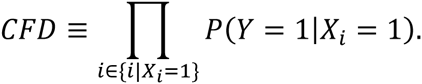

In contrast, a Naïve Bayes model would compute the probability that a guide-target pair is active given the feature values as

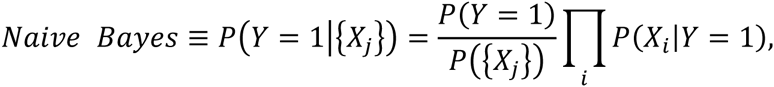

which makes only one assumption, namely, that conditioned on a guide being active, the features *X*_*i*_ are independent so that *P*({*X*_*j*_}|*Y* = 1)= Π_*i*_ *P*(*X*_*i*_|*Y* = 1). Using Bayes rule one can re-write the Naïve Bayes classifier as,

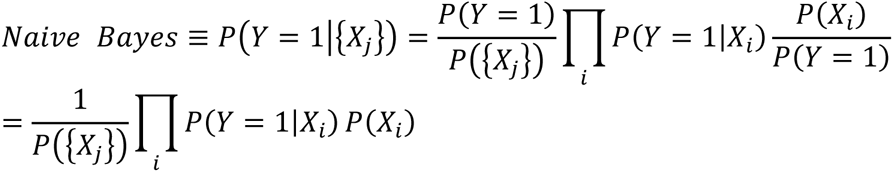

If we make two further assumptions, we find that Naïve Bayes classifier exactly matches *CFD*. The first assumption is that the features are marginally independent, namely that Σ_*i*_ *P*(*X*_*i*_) = *p*({*X*_*j*_}), in which case Naïve Bayes simplifies to

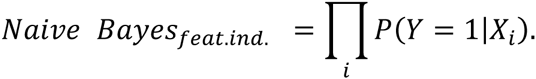

If we additionally assume that *P*(*Y* = 1|*X*_*i*_ = 0) = 1, then CFD and Naïve Bayes become identical. The CFD assumption of marginal feature independence seems reasonable and yielded good results. Consequently, we make the same assumption in our Elevation-score model. The second CFD assumption (*P*(*Y* = 1|*X*_*i*_ = 0) = 1) seems a more difficult one to accept, but with some careful thought (and the fact that CFD performs so well), also seems reasonable as we explain next; hence also make this assumption. The key insight is to ask which properties of the training data one expects to generalize to unseen data sets where the model might be applied. In particular, it seems reasonable to assume that (*P*(*Y* = 1|*X*_*i*_ = 1) is a quantity that will generalize to other data sets; intuitively the quantity reflects how likely a guide is to be active given that we observed a particular kind of mismatch (or alternate PAM)—as such, it is independent of the distribution of the types of mismatch (or alternate PAM) in the training versus test data sets. In contrast, *P*(*Y* = 1|*X*_*i*_ = 0) defines how likely a guide is to be active given that we did not observe a feature. When computing this quantity, one marginalizes over all examples in the CD33 data set where *X*_*i*_ = 0, which includes all guide-target pairs for which *X*_*j≠i*_ = 1; as such, this probability specifically depends on the distribution of mismatch/PAM types in the off-target data set, and their corresponding activity layers. Therefore, we don’t necessarily expect the quantity *P*(*Y* = 1|*X*_*i*_ = 0) to generalize from our training data (CD33-specific) to general test sets. Now the question remains, how can we therefore make a reasonable approximation? One could try to posit a canonical theoretical or actual data set which will best generalize; however, it is extremely difficult to come up with such a set. Furthermore, in light of how we are going to use our Naïve Bayes probabilities (described next), getting it exactly right is not critical. Hence we make the CFD assumption that *P*(*Y* = 1|*X*_*i*_ = 0) = 1.

We have now shown that with two assumptions, the *CFD* model can be interpreted as a Naïve Bayes classifier. The reason for making this connection is not only to put the *CFD* in a proper probabilistic framework, including its assumptions, but more importantly, to then generalize this model in order to improve its performance, which we now do in describing our Elevation-score model.

#### Elevation-score as two-layer stacked regression

We generalized away from CFD in three main ways: (i) we moved from classification to regression, (ii) we augmented the feature space, and (iii) we replace the *a priori* manner of combining by multiplication to combining using machine learning. We call the model implementing only the first two, Elevation-naïve, while we refer to the model resulting from all three as Elevation-score. We now explain these in more details.

The first observation in generalizing away from *CFD* is that it is a classification algorithm, which means it discards the real-valued assay measurements, converting them to be binary active/in-active. Thus, by design *CFD* is unable to capture the more nuanced information available in the data. In moving from classification to regression, the model has access to more fine-grained information. Although not widely used, there exist generalizations of Naïve Bayes classification to regression^25^; however, due to the specifics of our problem, they are not convenient to apply. Thus we developed our own approach in which we first convert the CD33 log-fold-change (LFC) values to lie in the range [0,1] so that they can, loosely-speaking, be interpreted as probabilities. To do so, we used a kernel density estimator to transform each LFC to the cumulative density of that LFC in the kernel density estimate. We used a Gaussian kernel and choose the bandwidth by 10-fold cross-validation (yielding 0.23).

Recall that Elevation-score is a two-layer stacked regression model where the first layer makes predictions for guide-target pairs with only a single mismatch (or alternate PAM), while the second-layer combines these for guide-target pairs with multiple mismatches (and alternate PAMs).

To learn each first-layer (single-mismatch) regression model, *P*(*y*|{*X*_*j*_}), we used Boosted regression trees (using default settings in scikit-learn) on the CD33 data.^26^ Since each guide-target pair has only a single *X*_*i*_ = 1 in these data, we could just have well used a linear regression model. However, we wanted to include a richer featurization of the guide-target pair than just features of the form A:G, 5, such that even for single mismatch data, more than one *X*_*i*_ = 1 (and some features were numeric rather than 0/1). Further, we wanted these features to be able to interact in a non-linear manner. Thus in addition to the CFD features, we also used “decoupled” versions of them—one of the form “A:G”, which was one-hot encoded (described at the end of this section) and the other an integer feature for the position (*e.g.*, 5). We also included whether the mutation was a transversion or a transition. We call the model which uses these improvements Elevation-naïve (it combines each mismatch/alternate PAM) just as CFD does, by multiplying the values together. As can be seen in Supplementary Figure 2, moving from classification to regression improved the performance of the off-target model, as did changing the features. Next we describe how we improve Elevation-naïve to obtain Elevation-score.

Although Elevation-naive improved upon *CFD*, there were several aspects of the modelling approach which suggested areas for further improvement. The first was that the Naïve Bayes assumption of class-conditional independence may not be fully justified. The second is that our regression model predicted values are not truly calibrated probabilities of guide-target activity; hence when we combine them under that assumption (as does Naïve Bayes and CFD), we may suffer in performance. Thus it stands to reason that if we could somehow loosen these assumptions, we might achieve better performance still. One way to do this is to augment the model, here with a second-layer, and then to use the limited amount of multiple-mismatch/PAM guide-target pair data to learn the newly added parameters. We refer to this second-layer of Elevation-score as the combiner because it learns how to combine the predictions from the single-mismatch model in a more nuanced way than simply multiplying them together like Elevation-naïve and CFD, thereby, allowing some of the stated assumptions to be mitigated. Thus where a *CFD*/Naïve Bayes approach would simply multiply single-mismatch probabilities together, we instead use a data-driven machine learning approach to fine-tune how they should be combined.

In particular, we first use our first-layer Boosted Regression trees model *J* times to make predictions for each of the *J* single mismatches/ alternative PAM (*i.e*., *J* features for which *X*_*j*_ = 1), yielding *J* predictions *ŷ*_*J*_ ∈ [0,1] (one for each feature with *X*_*j*_ = 1, and setting *ŷ*_*k*_ = 1 for the remaining *K* features that have *X*_*k*_ = 0). Therefore, each guide-target pair has *T* = *J* + *K* = 21 Boosted regression tree predictions {*ŷ*_*t*_} (20 for each possible mismatch position, and one for an alternate PAM). The log of 21 features ({log (*ŷ*_*t*_)}), along with their sum, their product, and *J*—the number of mismatches/alternate PAM are then the input to an L1-regularized linear regression combiner model—the second-layer model. We used each of the GUIDE-Seq data and the Hauessler *et al.* data to train a model, each time testing on the other data set. The final, deployed model used the one trained on the GUIDE-Seq data with 10-fold cross-validation to choose the regularization parameter. (When training on Hauessler *et al.* for Figure 2, the same settings were used). Note that Elevation-score’s two-layer model is inherently different from both a two-layer neural network^27^ and from stacked generalization^28^ because the data used to train each Elevation-score layer are different (single- *vs.* multi-mismatch).

Finally, because what we ultimately want are predictions of the probability that a guide-target pair is active, we also apply one final transformation to the output from the L1-regression model; namely, we then compute *p*(active|GUIDE-seq) using a logistic-regression model trained on the GUIDE-seq predictions (for the CD33 data using *Elevation-naive*) as inputs and using the binarized observed activities^1^ (LFC>1) as the target variable. Note that this transformation is monotonic and as such only affects performance of aggregation, not guide-target scoring, given that we use a Spearman rank correlation. Because for the aggregation task, the Spearman correlation is computed only after aggregation of scores for a guide, any change in scale of the pre-aggregated scores can dramatically influence the quality of the final aggregation, including a simple linear transformation.

#### One-hot encoding of categorical variables

A “one-hot” encoding refers to taking a single categorical variable and converting it to more variables each of which can take on the value 0, or 1, with at most, one of them being “hot”, or on. For example, with a categorical nucleotide feature which can take on values A/C/T/G, each letter would get converted into a vector of length four, with only one entry equal to on, corresponding to one of the four letters.

### Elevation-aggregate

In a sense, Elevation-score provides only the starting ingredients for choosing a guide with least expected off-target activity. To actually rank guides one needs to coalesce the scores from all guide-target pairs for a given guide into a single number so that guides can be ranked by this number for off-target activity. Thus we developed Elevation-aggregate to perform this task. Elevation-aggregate is based on gradient-boosted regression trees (using default settings in scikit-learn). The input features for the model were computed from the distribution of guide-target Elevation-score predictions and comprised: the mean, median, variance, standard deviation, 99^th^, 95^th^, 90^th^ percentiles, and sum of off-target scores. We compute these for each of: all off-targets, only genic off-targets, and only non-genic targets. Additionally, we compute these further features: sum of genic [non-genic] off-targets divided by total number of off-targets; fraction of targets that are genic; fraction that are non-genic; ratio of number of genic to non-genic targets; ratio of average genic to non-genic score. We found that adding these same features from the CFD model further boosted performance and so also included these. The final deployed model was trained only on the Avana data (combining with Gecko did not increase cross-validation performance).

### Comparison to other approaches

We compared models using the Spearman correlation between predicted and measured off-target activity. Furthermore, as discussed in the main text, we additionally evaluated the weighted-Spearman correlation for various weight settings in order to account for an asymmetrical loss with respect to false-positive-active errors as compared to false-negative-active errors. Specifically, we set the weights as follows. Let {*g*_*i*_} be the values of the normalized GUIDE-Seq or Hauessler data (all lying in [0, 1]). Then we set the weight for each data point to be *g*_*i*_ + *w* where *w* is varied through 10^−5^ to 100 and is denoted on the horizontal axis of the relevant figures. When *w* = 10^−5^, the weight is effectively equal to the GUIDE-Seq/Hauessler measured values. When *w* = 100, the weights become effectively identical for all guides, yielding a standard Spearman correlation.

For CFD we used the table provided by Doench et al. For CCTOP we re-implemented based on the description in their paper. For Hsu-Zhang guide-target pair scoring we re-implemented the approach based on the equation in their paper.

To compare Elevation to the aggregation scores of the MIT web server, each guide sequence was submitted to the MIT CRISPR Design Tool using their RESTful API provided for single sequences (http://crispr.mit.edu/). Every sequence was queried using sequence type “other region (23-500nt)” and target genome “human (hg19)” to obtain an off-target score. The server failed to produce scores for the following three sequences which we therefore removed from consideration in our comparison: sequence TGACCTGTGACCATGATCACCACAGGGTTG from Avana and sequences CAAGCCTGTGTGCTGCAAGCCTGTCTGCTCTGTGCC and TCTCTGGCCATCATTTCCTGGGAGAGATGGATGGTG from Gecko. All queries were submitted and their results processed between the dates of August 15^th^-29^th^ 2016, inclusive. No software version number was found in the output or web page.

To compare Elevation to the CFD server (http://portals.broadinstitute.org/gpp/public/analysis-tools/sgrna-design), each gene corresponding to an Avana or Gecko guide was submitted to between September 21^st^-23^rd^ 2016, inclusive, and the relevant guide rows retrieved. A score for a guide was obtained by adding the values in the two fields “Tier I Match Bin I Matches” and Tier I Match Bin II Matches” as done by Doench *et al.*^1^ Although the server returns a field off-target rank, this field cannot be readily compared across guides.

### Elevation-search

To perform efficient genomic searches for potential off-targets we developed the program dsNickFury which uses seed and extension^29^ (using two tandem seeds) to find near-match CRISPR-Cas9 targets. In brief, dsNickFury can leverage distributed computing to efficiently catalog every potential CRISPR-Cas9 target in a genome for any CRISPR-Cas9 system with targets that can be abstracted into some maximum length of RNA guide followed by a set of potential PAM sequences of fixed length. These potential targets are then organized into a tree data structure based upon two tandem seed sequences (lengths of 5 nucleotides used here, but this is a user-specified parameter) taken from the guide sequences immediately proximal to the PAM site. The first branch layer of the tree structure is comprised of all observed first tandem seeds (most proximal to the PAM) while the second layer contains branches for each second tandem seed. Each first and second seed combination links to a file containing all of the potential CRISPR-Cas9 targets in the genome that have that specific combination of tandem seeds proximal to the PAM site.

Potential off-target matches are defined by a maximum number of mismatches relative to the intended target (set by the user) with a certain number of bases distal to the PAM being ignored if desired. Here, mismatch tolerance was set to 3, with the 3 most distal bases from the PAM being ignored. All PAMs deemed to have non-zero activity in ref. ^1^ are considered (NAG, NCG, NGA, NGC, NGG, NGT, NTG), with non-NGG PAMs incurring a single mismatch count. This strategy was based upon previous observations that much CRISPR-Cas9 off-target activity risk is determined by the number of mismatches between on- and off-target sequences with bases more distal to the PAM sequence being more mismatch tolerant and contributing less to specificity.^3^ Potential sites are searched initially based upon their tandem seeds, using a depth-first search of the cached tree structure. Any leaves with fewer mismatches than the maximum allowed have the same check then applied to their extended sequences, ignoring bases distal to the PAM as determined by user-specified parameter. Those sequences that pass the filters are considered as potential off-targets and are scored by Elevation. These sites can be sorted based on their mismatch counts and/or Elevation score and in general can be reported directly to the user by way of a file or by deposition in to a NoSQL database as we have done here. We additionally use the Ensembl database to determine if each off-target is in an annotated gene or not, such that users can obtain an aggregated off-target score across one, the other, or both.

Because most sites can be disqualified based upon their seeds without loading the extended sequence and have already been annotated by both sequence and locus, searches can be conducted using significantly fewer resources than an alignment-based search. This allows for many searches to be conducted in parallel on a distributed computing environment. For results reported herein we pre-computed and stored all human genome-wide results for both on- and off-target predicted activities in a cloud-based database which we make available to the community.

Our system is designed to function on several different CRISPR-Cas9 systems with PAM sites at the 3’ end of the target. Parameters may be set for different lengths of guide sequence, PAM sequences with higher activity, and species of origin for the reference genome. Potential targets can be ranked for on-target efficiency and off-target risk. The system is currently using Azimuth for on-target activity prediction, and Elevation for off-target activity prediction for the *S. pyogenes* CRISPR-Cas9 system.

A summary of the search parameters used for all experiments in this paper as well as the on-line cloud service are as follows: we included all off-targets in the genome with no more than 3 mismatches in the 4-20 of the guide, with any number of mismatches in the first three guide nucleotides, and considering any PAM deemed to have non-zero probability according to the CFD model (namely, NAG, NCG, NGA, NGC, NGG, NGT, NTG).^1^ We stopped any searches that yielded more than 40,000 potential off-targets according to these criteria. For those yielding more than 40,000, we set our Elevation guide potential (i.e., the final guide aggregate value) to be equal to 1,000.

### Code Availability

All source code and a front-end website for the cloud service will be made available from http://research.microsoft.com/en-us/projects/crispr upon publication.

## Acknowledgments

We thank Akshaya Annavajhala for Azure cloud support, Carl Kadie for use of his HPC cluster code, Jim Jernigan, Oleg Losinets and the HPC team for cluster support, Mudra Hegde for help with data, and Maximilian Haeussler for help accessing data from his paper. Michael Weinstein is supported by a UCLA Collaboratory Fellowship. This work used computational and storage services associated with the Hoffman2 Shared Cluster provided by UCLA Institute for Digital Research and Education’s Research Technology Group, and made use of an Azure-for-Research grant to UCLA.

## Author Contributions

JL and NF designed, implemented and evaluated the machine learning and statistical methods (Elevation-score, Elevation-aggregate, Elevation). MW designed, implemented and ran the Elevation-search infrastructure, also known as dsNickFury. JD provided biological expertise relevant to all parts of the work. LH created a front end webpage for the cloud service. ME ran and implemented experiments. JL, MW, JD, NF wrote the paper.

## Supplemental Figures

**Supplemental Figure 1.**
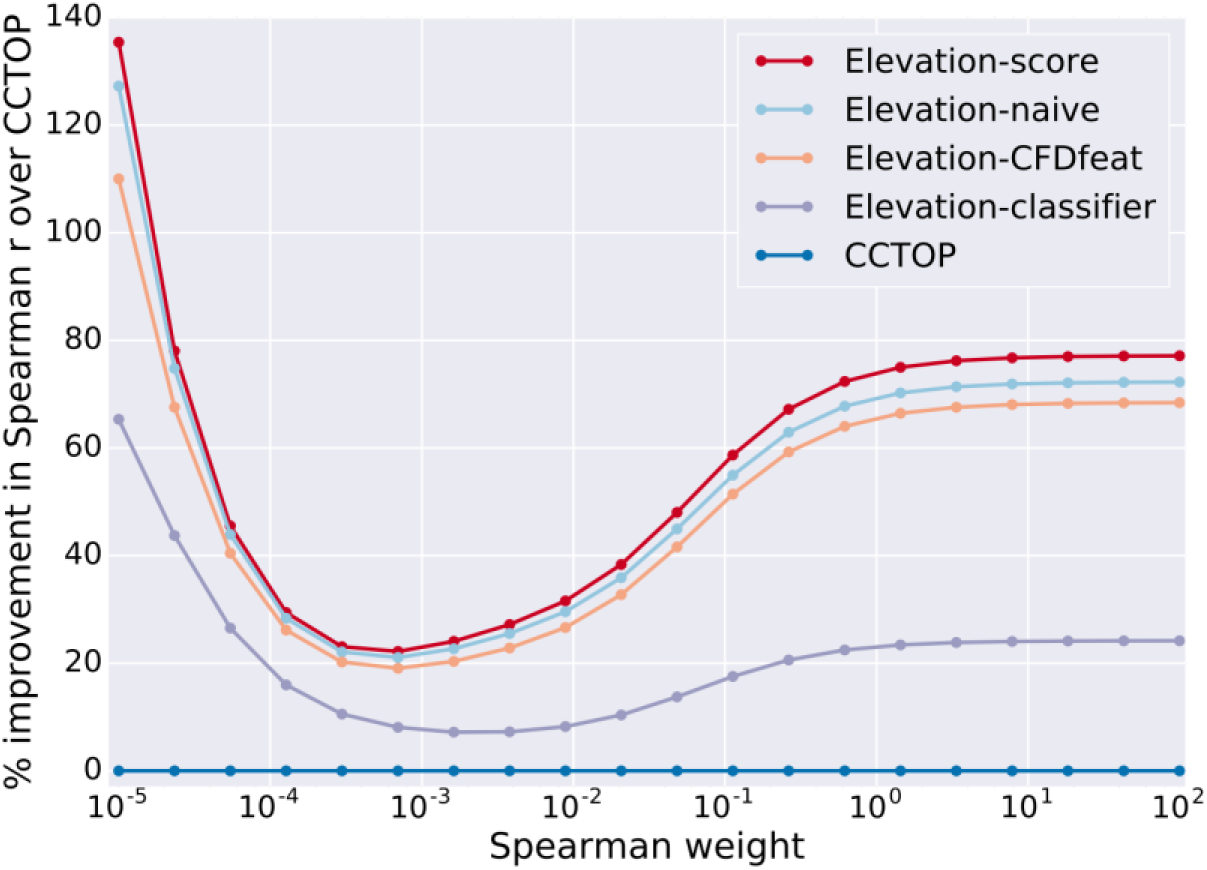
Impact of modelling decisions on guide-target scoring. Decrease in performance owing to skipping the second layer model, instead using the product of all outputs from the first-layer model (Elevation-naïve); using only features corresponding to those in CFD (the joint position and mismatched nucleotide identities) in Elevation-naive (Elevation-CFDfeat); using classification instead of regression in Elevation-naïve (Elevation-classifier). Note that the actual CFD model itself if plotted would exactly overlay Elevation-classifier. The percent improvement over a baseline of CCTOP is used for consistency with plots in the main paper. Results shown are for training on GUIDE-Seq and testing on Haeussler *et al.*

